# Non-viral *PDCD1* locus-directed hYP218 CAR T cells with enhanced anti-tumor activity against mesothelin-positive solid tumors

**DOI:** 10.64898/2026.11.18.767645

**Authors:** Sameer Mir, Abhilash Venugopalan, Raj Chari, Yosef Charkatli, Jingli Zhang, Qun Jiang, Raffit Hassan

## Abstract

Chimeric antigen receptor (CAR) T-cell therapy has shown limited efficacy against solid tumors partly due to T-cell dysfunction and the immunosuppressive tumor microenvironment. Precise, non-viral genome engineering provides an opportunity to generate CAR T cells with defined genomic integration while simultaneously introducing genetic modifications that enhance T-cell function. Here, we developed a CRISPR/Cas9-mediated homology-directed repair (HDR) strategy to integrate a mesothelin-targeted hYP218 CAR into either the endogenous *PDCD1* (PD1) locus or the *AAVS1* safe-harbor locus and compared these products with conventionally generated lentiviral CAR T cells. Targeted integration into the PD1 locus generated functional hYP218 CAR T cells with complete abrogation of PD1 expression and potent antigen-specific cytotoxicity against mesothelin-positive tumor cells. Despite exhibiting substantially lower CAR surface expression, PD1-site integrated CAR T cells consistently outperformed AAVS1-directed CAR T cells and achieved antitumor activity comparable to lentiviral CAR T cells. Following repetitive antigen stimulation, PD1-site integrated CAR T cells maintained cytotoxic function, exhibited reduced expression of exhaustion-associated markers, and preserved a less differentiated phenotype. Compared with lentiviral CAR T cells, non-viral CAR T-cell products secreted lower levels of IFN-γ and TNF-α while maintaining robust antitumor activity, consistent with more controlled effector activation. Collectively, these findings establish targeted integration of the hYP218 CAR into the endogenous *PDCD1* locus as an effective strategy for generating potent mesothelin-directed CAR T cells for treatment of solid tumors.

## Introduction

CAR T cell therapy has revolutionized cancer immunotherapy, offering durable remissions for patients with relapsed or refractory hematologic malignancies, including B-cell leukemias and lymphomas^1–4^. However, the successful application of CAR T cell therapy to solid tumors remains challenging, primarily due to the immunosuppressive tumor microenvironment, antigen heterogeneity, and practical limitations associated with conventional viral vector-based manufacturing^5–7^. Traditional CAR T cell production relies on retroviral or lentiviral vectors for gene delivery, which despite their efficacy have some drawbacks, including high manufacturing costs, extended production timelines, unpredictable transgene integration, and the risk of insertional mutagenesis^8–11^. Furthermore, lentivirally engineered CAR T cells often exhibit excessive cytokine release and tonic signaling, leading to premature T cell exhaustion and diminished antitumor activity in solid tumor settings^12–14^. These limitations highlight the need for more scalable, predictable, and biologically compatible strategies to generate potent and persistent CAR T cells.

Mesothelin (MSLN) is an attractive target for CAR T-cell therapy because it is highly expressed in a wide range of solid malignancies, including mesothelioma, pancreatic adenocarcinoma, ovarian cancer, gastric cancer, and other epithelial tumors, while exhibiting limited expression in normal tissues^15–17^. However, results of anti-mesothelin CAR T cell clinical trials have shown limited activity in patients^18–20^. One potential reason for this lack of activity is due to the antibody fragment used to make the CAR T cells binding to shed mesothelin in serum or tumor microenvironment and limiting the efficacy of CAR T cells^21^. We have recently shown that hYP218 CAR T cells that recognizes a membrane-proximal epitope of mesothelin has increased antitumor activity, tumor infiltration, and persistence in preclinical studies^22,23^. Nevertheless, the efficacy of mesothelin-targeted CAR T cells may also be constrained by the immunosuppressive tumor microenvironment and progressive T-cell dysfunction associated with chronic antigen exposure.

Programmed cell death protein 1 (PD1) is a major inhibitory immune checkpoint that contributes to T-cell exhaustion in solid tumors^24^. Previous studies have demonstrated that PD1 disruption or blockade can improve CAR T-cell function and antitumor activity^25,26^. In parallel, advances in CRISPR/Cas9-mediated genome engineering have enabled site-specific integration of CAR transgenes into defined genomic loci, thereby reducing the risks associated with random viral integration and providing greater control over transgene expression^27,28^. Several groups have reported non-viral CAR T-cell engineering approaches, PD1-disrupted CAR T cells, and site-specific CAR integration strategies^28,29^. However, the therapeutic potential of combining a clinically relevant mesothelin-targeted hYP218 CAR with precise non-viral integration into the endogenous PD1 locus has not been investigated in solid tumors.

Targeting the CAR transgene to the *PDCD1* locus enables simultaneous CAR integration at a defined genomic site and disruption of PD1-mediated inhibitory signaling, simplifying the manufacturing workflow compared with sequential PD1 knockout and CAR delivery. To establish a non-viral CAR T-cell engineering platform for mesothelin-expressing solid tumors, we engineered primary human T cells with the clinically relevant hYP218 CAR using a CRISPR/Cas9-based approach. The CAR transgene was targeted to either the *PDCD1* locus or a safe-harbor locus to evaluate the impact of integration site on CAR T-cell phenotype and function. We further compared these non-viral hYP218 CAR T-cell products with conventional lentivirally transduced hYP218 CAR T cells, evaluating CAR expression, T-cell expansion, cytokine production, differentiation and exhaustion-associated phenotypes, and antitumor activity against solid tumors.

## Materials and methods

### Cell lines

Established tumor cell line OVCAR-8 and KLM-1 were gifts from Dr. Hisataka Kobayashi and Dr. Christine Alewine from NIH (Bethesda, MD) respectively. The early-passage mesothelioma cell line NCI-Meso63 was established from ascites or pleural fluid obtained from a patient with mesothelioma treated at the NCI (Bethesda, MD) under an Institutional Review Board (IRB) approved protocol (ClinicalTrials.gov NCT 01950572). The methods for establishing primary cell culture lines have been described previously^30^. All the cells were maintained in RPMI1640 medium (Gibco) with 10% FBS (GemCell) 2 mmol/L L-glutamine, and 100 U penicillin– streptomycin (Invitrogen), in a humidified atmosphere containing 5% CO_2_ at 37°C. These cell lines were authenticated using short tandem repeat profiling and regularly tested for the absence of Mycoplasma. For *in vivo* bioluminescence detection, all cell lines were modified to express luciferase (Luc^+)^ and enhanced green fluorescent protein (eGFP^+^).

### T cell electroporation and transduction

Electroporation of T cells was performed using an optimized protocol, ensuring high transfection efficiency and cell viability. Post-electroporation, cells were cultured in AIM-V medium (Gibco) supplemented with IL-2, IL-7, and IL-15 (PeproTech), and CAR expression was confirmed by flow cytometry (Beckman Coulter). Electroporation was performed 48 hours after T cell stimulation using a Lonza 4D electroporation system (Lonza) with Program EO115. Lentivirally transduced hYP218 CAR T cells were generated as previously described^22^. VSVG pseudotyped lentivirus with a high titer was developed by Cellomics Technology (Halethorpe, MD). Detailed methods for construction of the hYP218 CAR cassette, ribonucleoprotein and linear dsDNA production, PBMC isolation and T cell electroporation are provided in the Supplementary Information.

### Flow cytometry and CAR characterization

CAR expression and T-cell surface phenotypes were assessed by flow cytometry (CytoFlex; Beckman Coulter) and analyzed using FlowJo (BD Biosciences; version 10.8.1). hYP218 CAR expression was determined by staining for the co-expressed truncated human EGFR (huEGFRt), and CAR-positive cells were quantified within the CD3 population. PD1 expression was evaluated in CD3 CAR cells and compared among LV-hYP218, AAVS1-hYP218, and PD1-hYP218 CAR T cells. Estimated CAR surface density was quantified using BD Quantibrite™ PE beads and PE-conjugated anti-human EGFR staining. Because huEGFRt is co-expressed with the CAR cassette through a T2A sequence, huEGFRt-derived PE molecules per cell were used as an estimate of relative CAR surface density. Detailed staining procedures, Quantibrite calibration, and antibodies used for flow cytometry are provided in the Supplementary Information and Table S1.

### *In vitro* functional assays

Cytotoxicity and cytokine production were evaluated following coculture of CAR T cells with MSLN-expressing OVCAR-8, KLM-1, or NCI-Meso63 cells. Tumor-cell viability was quantified using luciferase-based assays, and IFN-γ and TNF-α secretion was measured by ELISA. For repeated antigen stimulation, CAR T cells were cocultured with OVCAR-8 cells over 7 days, with serial assessment of cytotoxicity and T-cell phenotype. Equivalent numbers of CAR-positive T cells were used across CAR T-cell products. Detailed experimental procedures are provided in the Supplementary Information.

### *In vivo* tumor models

All animal experiments were conducted in compliance with the protocol approved by the Animal Care and Use Committee (ACUC) of the NCI, NIH. NOD SCID gamma (NSG) mice were obtained from The Jackson Laboratory (Bar Harbor, ME, USA). All mice were housed in a pathogen-free environment following NIH guidelines approved by the NCI’s ACUC. For the experiment comparing CAR T groups in the OVCAR-8 and NCI-Meso63 models, 5- to 6-week-old female NSG mice were intraperitoneally injected with 5 × 10^6^ luciferase-expressing OVCAR-8 and NCI-Meso63 cancer cells and allowed to grow for 7 days. Subsequently, a single dose of 3 × 10^6^ CAR T cells was administered intravenously. For the experiment comparing CAR T groups in the highly aggressive KLM-1 model, 5- to 6-week-old female NSG mice were intraperitoneally injected with 0.5 × 10^6^ luciferase-expressing KLM-1 cancer cells and 7 days after tumor implantation, were administered a single dose of 3 × 10^6^ CAR T cells. For all in vivo experiments, the indicated doses refer to CAR-positive T cells. The frequency of CAR-expressing cells in each product was determined by flow cytometry, and the total number of infused cells was adjusted to ensure administration of equivalent numbers of CAR-positive T cells across the different treatment groups. Bioluminescence images were acquired and analyzed using the IVIS Imaging System and software (PerkinElmer, IVIS Lumina *III In vivo* Imaging System) after intraperitoneal injection of the substrate D-luciferin (Syd Labs, Cat# MB000102-R70170). Mice were randomized to different groups based on tumor radiance before CAR T cell injection. Mice were euthanized in accordance with the experimental protocols or when they met prespecified endpoints defined by the ACUC, NCI, NIH.

### Persistence and characterization of tumor-infiltrating hYP218 CAR T cells

To evaluate the anticancer activity of CAR T cells at low doses and the phenotypic status of cell surface markers of hYP218 CAR T cells, we used the NCI-Meso63primary mesothelioma NSG model. 5 × 10^6^ luciferase-expressing NCI-Meso63 cells were injected intraperitoneally into NSG mice and were allowed to grow for 14 days. Subsequently, 1 × 10^6^ CAR T cells were injected intravenously. Mice were IVIS imaged a day before termination of the experiment, and the tumors were harvested from the mice for comprehensive analysis of phenotypic markers on days 7 and 14 post-treatment. This allowed us to evaluate the presence and functional state of the infused hYP218 CAR T cells. Liberase TL enzyme (Roche), in conjunction with mechanical procedures, was used to produce single-cell suspensions from the extracted tumor tissues. Flow cytometry was used to assess the expression of various immunological markers, including those indicative of naive, activation, and exhaustion-associated phenotype.

### Statistical analysis

An unpaired Student’s t-test was used to compare differences among LV, AAVS1, and PD1-hYP218 CAR T cells. Statistical analysis was performed using GraphPad Prism 9.0. P < 0.05 was considered statistically significant.

## Results

### Generation and characterization of non-viral AAVS1 and PD1-hYP218 CAR T cells

To develop a non-viral platform for targeted hYP218 CAR T-cell generation, we used CRISPR/Cas9-mediated homology-directed repair (HDR) to target the hYP218 CAR cassette to either the PD1 or AAVS1 locus (Figure 1A). PD1 was selected as the primary knock-in site to combine CAR integration with disruption of an inhibitory immune checkpoint, whereas AAVS1 served as a safe-harbor control. We first optimized the donor DNA used for targeted CAR integration. The 7,072-bp PD1-hYP218 CAR plasmid was either linearized by single digestion or double-digested to remove bacterial propagation sequences, followed by purification of the 5,363-bp linear CAR donor fragment. Compared with the undigested plasmid and single-digested linear donor, the purified double-digested donor resulted in greater T-cell expansion and a higher percentage of CAR-positive T cells following nucleofection (Suppl Fig. 1A-B). Based on these results, the purified double-digested donor was used for subsequent HDR-mediated CAR integration. We observed a CAR expression of approximately 10% (Figures 1B and 1C) using linear double-stranded DNA (dsDNA) with 800-bp homology arms as the repair template for site-specific integration at the AAVS1 and PD1 loci. Post-electroporation, T cells fully recovered from electroporation-induced stress within 72 hours and showed robust expansion, achieving at least 20-fold expansion within a week (Figure 1D). Insertion of the hYP218 CAR cassette into the PD1 locus (PD1-hYP218 CAR T cells) resulted in a reduction of PD1 expression to 0.96%, compared to 6.49% in AAVS1-hYP218 CAR T cells and 16.9% in LV-hYP218 CAR T cells (Figures 1E and 1F).

**Figure 1.**
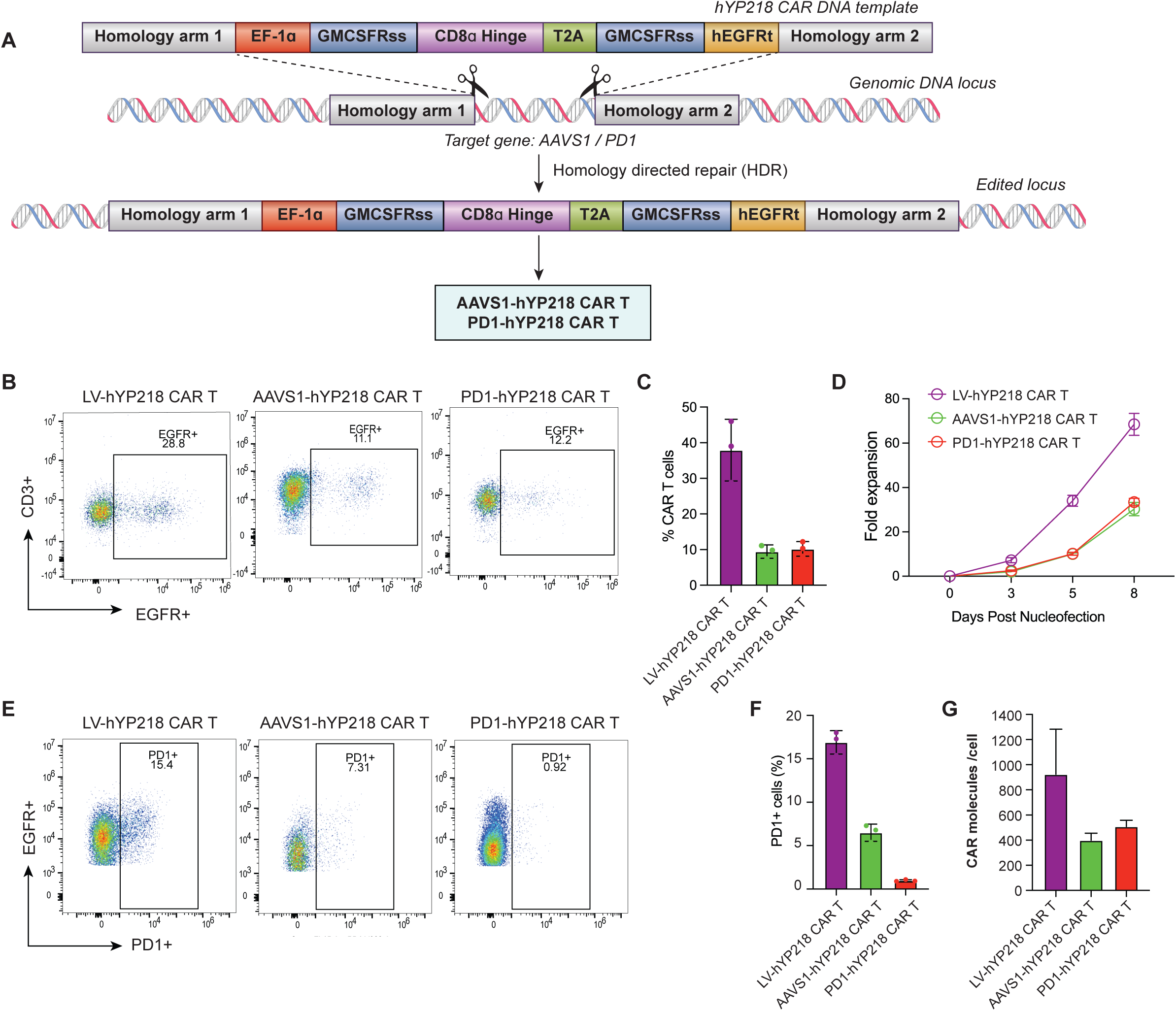
Generation and validation of non-viral hYP218 CAR T Cells. (A) Schematic representation of the electroporation workflow. The hYP218 CAR cassette was specifically integrated into the *AAVS1* or *PDCD1* locus via CRISPR/Cas9-mediated homologous recombination. (B-C) Representative example and summary of multiple experiments (n = 3) illustrating the percentage of hYP218 CAR-positive cells detected by flow cytometry 7 days post-electroporation using equimolar amounts of DNA templates. Lentivirus (LV) transduced hYP218 CAR T cells were used as control. (D) Fold expansion of the hYP218 CAR T cells following electroporation. (E-F) Representative example and summary of multiple experiments (n=3) showing efficiency of PD1 knockout in CD3^+^CAR+ cells. Flow cytometry was used to determine the percentage of PD1-expressing cells within the CD3+CAR+ gated population 7 days after electroporation. (G) CAR density of hYP218 CAR T cell products. Data are presented as mean ± SD from 3 independent experiments.

Furthermore, we quantified the estimated hYP218 CAR surface density using BD Quantibrite Quantitation assay. LV-hYP218 CAR T cells exhibited the highest CAR density, averaging 924 molecules per cell, followed by PD1-hYP218 CAR T cells with 508 and AAVS1-hYP218 CAR T cells with 399 molecules per cell (Figure 1G). Together, these data show that non-viral genome-engineering enables precise, locus-specific integration of hYP218 CAR into the T cell genome, with high viability and expansion potential and is associated with differences in surface CAR density that may contribute to the distinct functional profiles among the engineered CAR T-cell products.

### Non-viral hYP218 CAR T cells are cytotoxic to mesothelin expressing cell lines with reduced cytokine release

To assess cytotoxicity, AAVS1 and PD1-hYP218 CAR T cells were co-cultured with mesothelin expressing cancer cell lines OVCAR8 (ovarian cancer), NCI-Meso63 (mesothelioma), and KLM-1 (pancreatic cancer) at varying effector-to-target (E:T) ratios for 24 hours. Lentivirally transduced hYP218 CAR T cells (LV-hYP218) were included for comparison, and untransduced T cells served as negative controls. All three hYP218 CAR T-cell products exhibited robust cytotoxicity against the mesothelin-expressing tumor cell lines tested (Figure 2A). PD1-hYP218 CAR T cells demonstrated cytotoxic potency comparable to LV-hYP218 CAR T cells, as reflected by similar ET50 values across the tumor cell lines. In contrast, AAVS1-hYP218 CAR T cells generally exhibited higher ET50 values, indicating that a greater effector-to-target ratio was required to achieve equivalent tumor-cell killing. At an E ratio of 3:1, cytotoxicity was comparable among all three CAR T-cell products against OVCAR-8 and NCI-Meso63 cells, whereas differences among the CAR T-cell products were more apparent against KLM-1 cells. To evaluate functional responses and the risk of cytokine-mediated toxicities, we quantified effector cytokine release by hYP218 CAR T cell products using ELISA. Non-viral hYP218 CAR T cells secreted significantly lower levels of IFN-γ and TNF-α compared to LV-hYP218 CAR T cells during co-culture with OVCAR-8 and KLM-1 tumor cells (Figure 2B). This reduction in cytokine secretion can potentially decrease toxicity in patients, although additional studies incorporating broader cytokine analyses and clinically relevant models will be required to confirm these findings.

**Figure 2.**
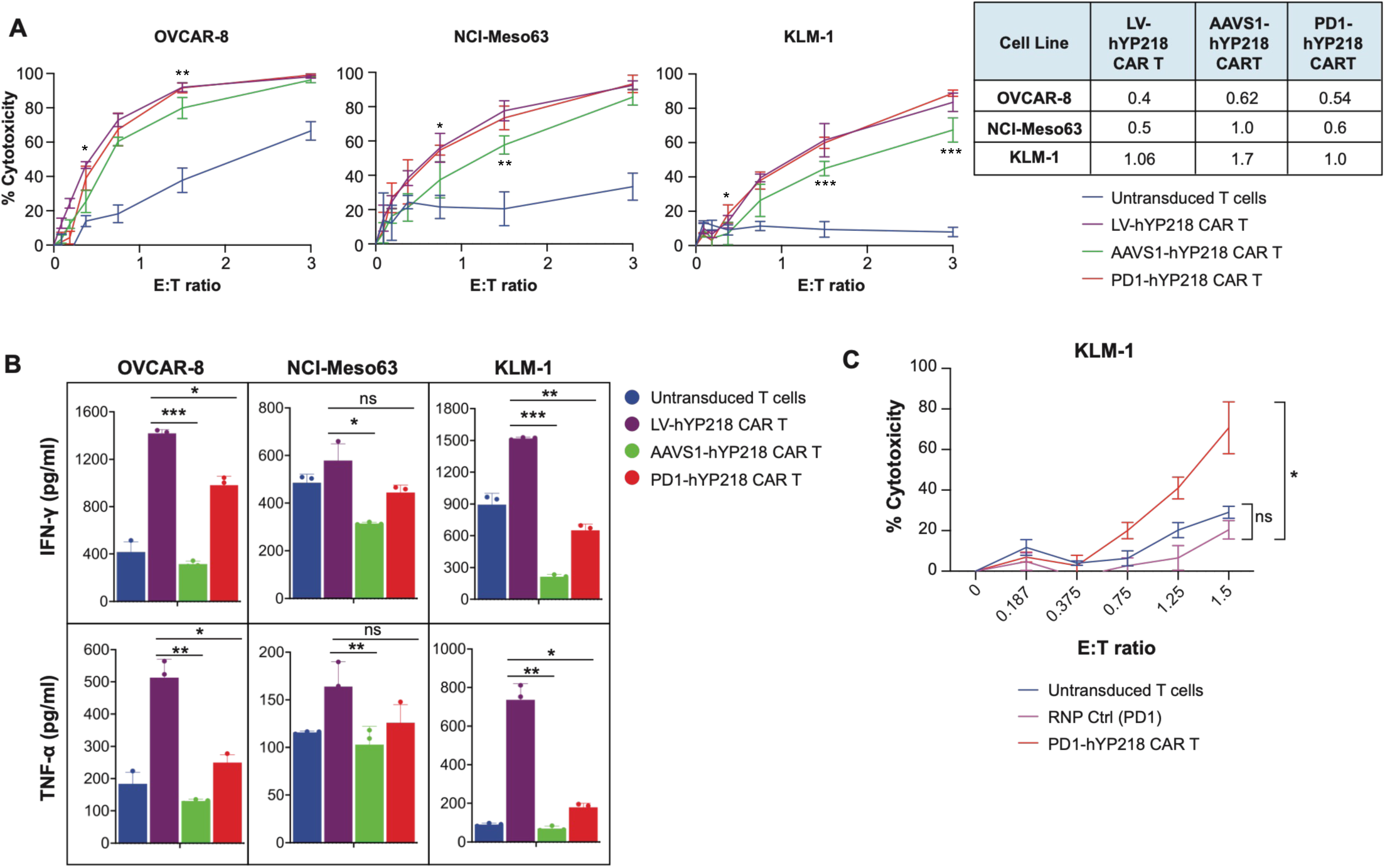
In vitro cytotoxicity of lentiviral and non-viral hYP218 CAR T cells. (A) In vitro cytotoxicity and percentage of tumor cell killing were assessed against mesothelin-positive OVCAR-8, NCI-Meso63, and KLM-1 tumor cell lines at varying E:T ratios using a luciferase assay. Statistical significance comparing groups PD1-hYP218 and AAVS1-hYP218 CAR T were shown. The mean value of ET50 was shown in the table. (B) ELISA was used to measure IFN-γ and TNF-α cytokine secretion in the supernatant after co-culture of hYP218 CAR T cells with mesothelin-expressing cancer cells for 24 hours. (C) The cytotoxicity against KLM-1 tumor cells by CAR T cells. Control samples were electroporated in the same manner as PD1-hYP218 CAR T cells, except that they lacked the hYP218 CAR DNA template. Data represent ± SD (n = 3). Significance levels: ns = non-significant; *p≤ 0.05; **p ≤ 0.01; ***p ≤ 0.001.

Additionally, editing the PD1 locus alone in T cells using a PD1-specific RNP produced basal cytotoxicity, similar to that of untransduced T cells. In contrast, electroporation with the hYP218 CAR gene induced robust cytotoxicity (Figure 2C), demonstrating that tumor cell killing is specifically attributable to hYP218 CAR expression and activity rather than to PD1 locus modification alone. Collectively, these findings indicate that non-viral PD1-hYP218 CAR T cells exhibit robust cytotoxicity against different tumor cell lines with reduced cytokine release.

### Non-Virally transduced hYP218 CAR T cells display sustained cytotoxicity following repetitive tumor cell challenge

CAR T cells in solid tumors face sustained antigen exposure and a suppressive microenvironment, which can decrease their activity over time. We therefore used a repetitive killing assay to model prolonged tumor antigen exposure and assess the durability of cytotoxic function, activation dynamics, and the development of exhaustion-associated phenotypes of the non-virally transduced hYP218 CAR T cells as compared with LV-hYP218 CAR T cells. The hYP218 CAR T cells were cocultured with OVCAR8 cancer cells at an E:T ratio of 3:1 and fresh OVCAR-8 cells were added on days 1, 3 and 5 (Figure 3A). We used basal RPMI medium supplemented with 10% FBS, without any added cytokines, including IL-2, thereby maintaining T cell proliferation and function solely through T cell interactions with cancer cells. Both lentiviral and non-virally transduced hYP218 CAR T cells displayed comparable levels of cytotoxicity after two rounds of restimulation; however, the level of cytotoxicity dropped upon third restimulation with OVCAR-8 cells (Figure 3B). We then investigated the phenotypic biomarker expression by different types of CAR T cells using flow cytometry (Figure 3C-G, Suppl Fig. 2A-E). All the hYP218 CAR T cell products exhibited a marked increase in CD69+ cells following the initial co-culture, followed by a decline over subsequent restimulations with OVCAR-8 cells (Figure 3C, Suppl Fig. 2A). In contrast, untransduced T cells maintained a consistently low level of CD69 expression throughout the time course, consistent with the absence of antigen-specific activation. Notably, the PD1 knockout CAR T cell construct displayed the highest level of CD69 expression on Day 1. Similarly, for the activation marker CD39, we observed a trajectory of increased expression after 24 hours of stimulation, followed by a decrease at subsequent time points (Figure 3D, Suppl Fig. 2B). LV and AAVS1-hYP218 CAR T cells exhibited early PD1 induction with a peak of PD1+ cells post 24-hour after co-culture, after which the levels returned to basal levels and remained stable until day 7. PD1+ cells peaked at nearly 60% in the LV-hYP218 CAR T cells, compared to approximately 40% in the AAVS1-hYP218 CAR T cells. As expected, PD1 expression was barely detected in PD1-hYP218 CAR T cells across repeated stimulations at all time points, given that CAR was inserted at the *PDCD1* locus (Figure 3E, Suppl Fig. 2C). A similar trend was observed in the expression levels of TIM3+ and LAG3+ by CAR T cells over time but there was no difference in lentivirus or non-virally transduced hYP218 CAR T cells (Figure 3F-G, Suppl Fig. 2D-E). Overall, these findings suggest that PD1-hYP218 CAR T cells show efficacy comparable to LV-hYP218 CAR T cells in a repetitive killing assay with similar expression of activation and exhaustion markers except PD1 since these cells lack it.

**Figure 3.**
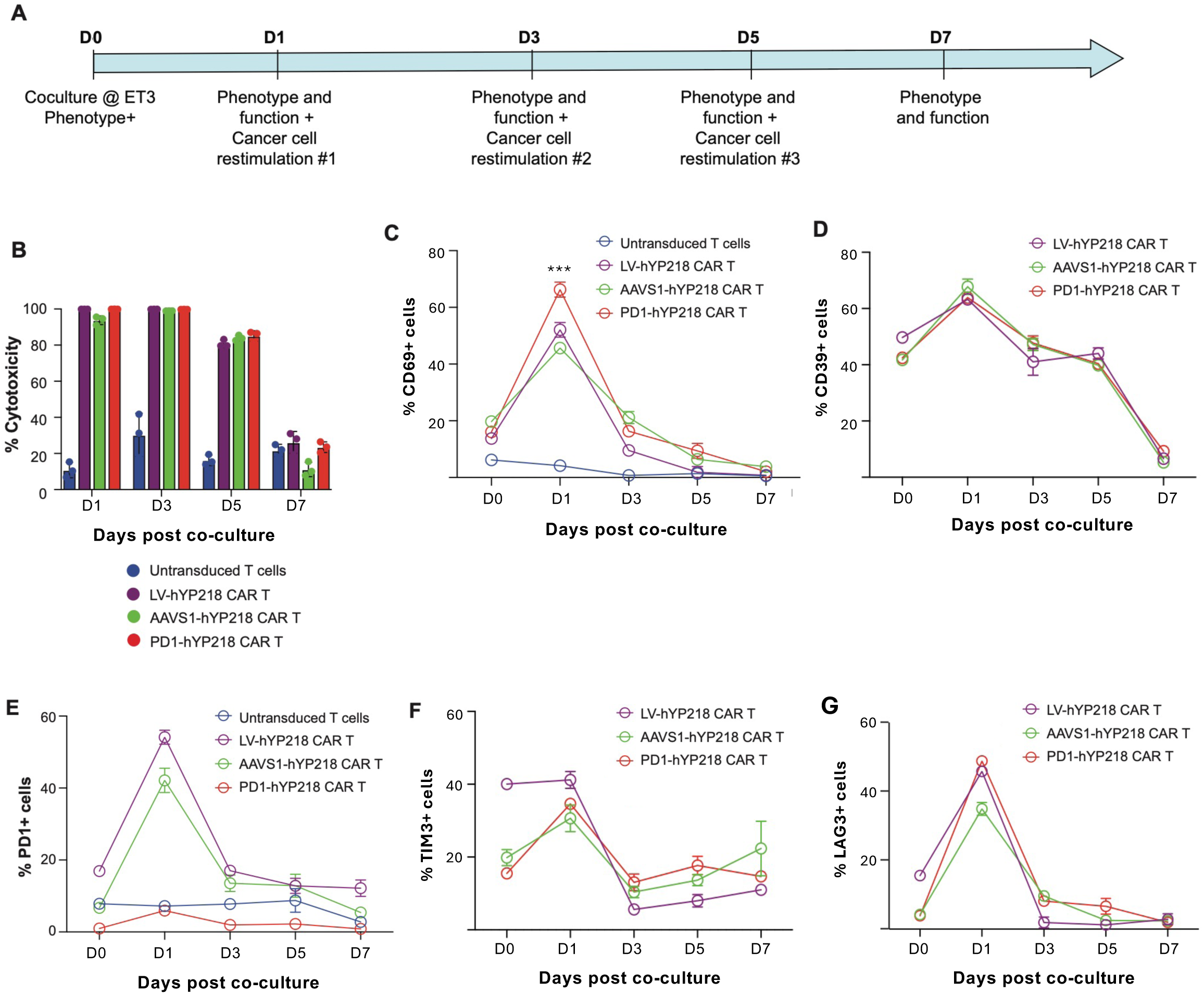
Phenotypic analysis of lentiviral and non-viral hYP218 CAR T cells following repeated co-culture with tumor cells. (A) Timeline of the experiment showing the specific days on which fresh tumor cells were added for re-stimulation and when the phenotypic analysis was done. Lentiviral and non-virally transduced hYP218 CAR T cells, along with untransduced T cell controls, were co-cultured with OVCAR-8 tumor cells for 7 days. Restimulation with fresh tumor cells was performed on days 1, 3, and 5. An effector-to-target (E:T) ratio of 3:1 was used, and CAR expression was normalized between groups before each stimulation. (B) Cytotoxicity of T cells against tumor cells at different timepoints. (C-G) The percentage of indicated biomarker positive CAR T cells was measured over multiple tumor cell re-stimulations to evaluate activation and exhaustion-associated profiles of lentiviral and non-virally transduced hYP218 CAR T cells. Data are plotted as mean ± SD (n = 3). Statistical significance comparing groups PD1-hYP218 and AAVS1-hYP218 CAR T were shown. *p≤ 0.05; **p ≤ 0.01; ***p ≤ 0.001.

### Non-Virally transduced PD1-hYP218 CAR T Cells exhibit efficacy in multiple tumor models

We then evaluated the efficacy of non-viral PD1-hYP218 CAR T cells against mesothelin-positive solid-tumor xenograft models in NSG mice, including OVCAR-8, NCI-Meso63, and KLM-1. NSG mice were injected with cancer cells (5 x 10^6^ for OVCAR-8 and NCI-meso63; 0.5 x 10^6^ for KLM-1) intraperitoneally and one week later after the tumors were established treatment was initiated with a single intravenous injection of 3 x 10^6^ hYP218 CAR T cells. Saline injection and non-transduced T cells were used as controls (Figure 4A). In the OVCAR-8 model, PD1-hYP218 CAR-T cells as well as LV-hYP218 CAR-T cells had the most significant impact on tumor growth and complete tumor clearance was achieved by week 2 in all the treated mice. By contrast, AAVS1-hYP218 CAR T cells resulted in tumor eradication in only 60% (3/5) of mice and this effect was delayed as compared to the other two CAR T cell treated groups (Figures 4B-C). In the NCI-Meso63 mesothelioma model, both PD1-hYP218 CAR T cells and LV-hYP218 CAR T cells achieved complete tumor clearance by week 4, whereas residual tumors remained in mice treated with AAVS1-hYP218 CAR T cells (Figures 4D-E). In the aggressive KLM-1 pancreatic cancer model, mice treated with PD1-hYP218, or AAVS1-hYP218 CAR T cells had significantly lower tumor burdens that was comparable to LV-hYP218 CAR T cells and markedly lower than those in mock- or saline-treated controls (Figures 4F-G). Collectively, these results demonstrate that non-virally transduced hYP218 CAR T cells, particularly PD1-hYP218 CAR T cells, exhibit potent anti-tumor activity similar to treatment with LV-hYP218 CAR T cells.

**Figure 4.**
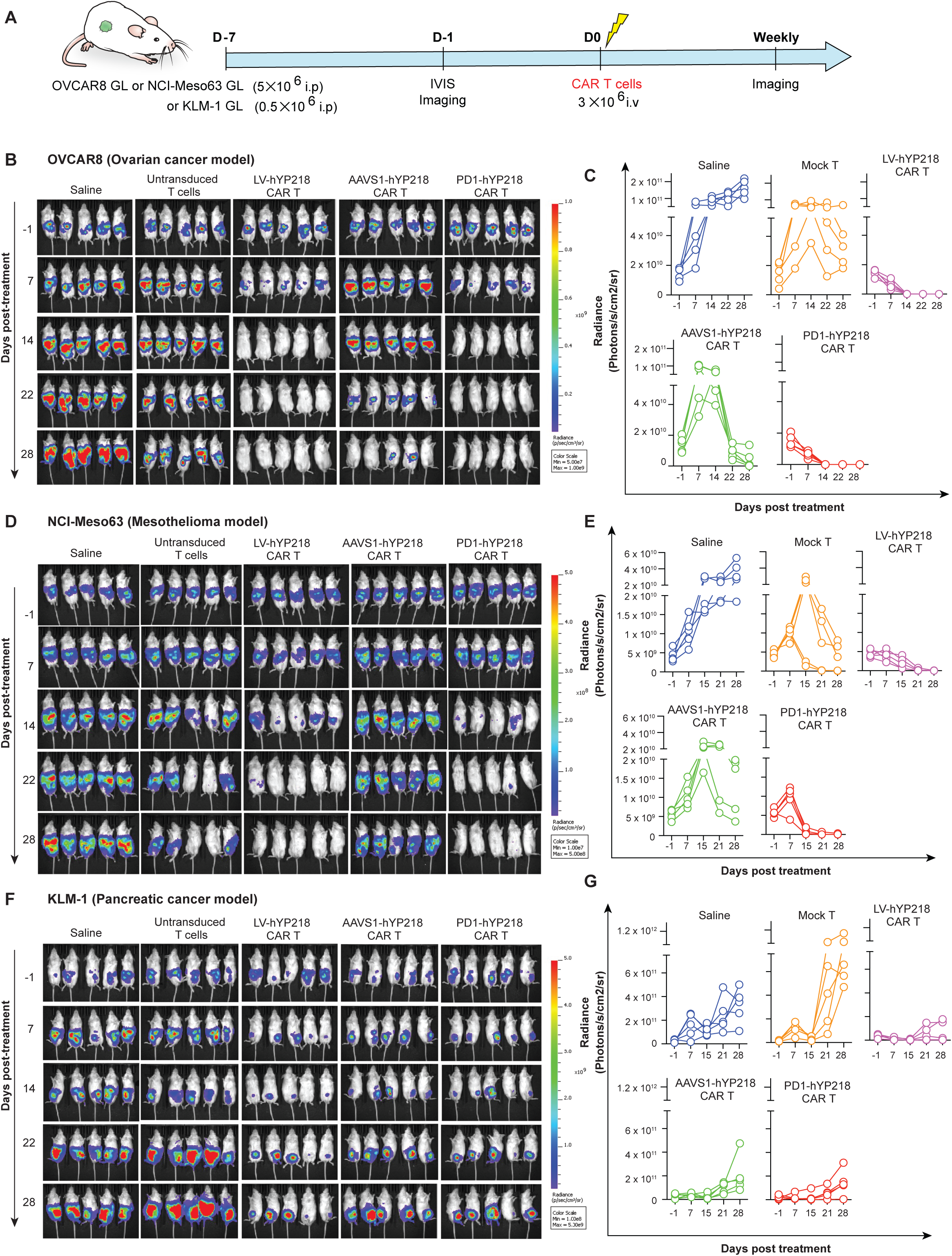
Anti-tumor efficacy of PD1-hYP218 CAR T Cells. (A) Experimental design for treating tumor bearing NSG mice (n=5 per treatment group) with lentiviral and non-virally transduced hYP218 CAR T cells. NSG mice were intraperitoneally injected with 5 × 10^6^ firefly luciferase (ffLuc)-transduced OVCAR-8, NCI-Meso63 or 0.5 x 10^6^ KLM-1tumor cells followed by intravenous administration of 3 × 10^6^ hYP218 CAR T cell products on day 7 post-tumor inoculation. Tumor burden was monitored by bioluminescence imaging. (B-C) Bioluminescence images and intensity showing tumor growth in the OVCAR-8 NSG tumor model following treatment with PD1-hYP218 or LV-hYP218 CAR T cells. (D-E) Bioluminescent monitoring and quantification in the primary NCI-Meso63 mesothelioma NSG model. (F-G) Tumor growth in the highly aggressive KLM-1 xenograft model treated by virally or non-virally transduced hYP218 CAR T cells, as measured by IVIS BLI imaging and quantification.

### PD1-hYP218 CAR T Cells do not upregulate PD-1 expression and maintain a naïve phenotype in the tumor microenvironment

To evaluate the phenotypic of PD1-hYP218 CAR T cells within the tumor, we used the luciferase-expressing patient-derived NCI-Meso63 xenograft model. NSG mice were injected intraperitoneally with the tumor cells and, after 2 weeks, were treated with a lower dose of 1 × 10^6^ hYP218 CAR T-cell products: LV-hYP218, AAVS1-hYP218 or PD-1 hYP218 CAR T cells; untransduced mock T cells were used as a control (Figure 5A). Tumor progression was monitored longitudinally using the IVIS bioluminescence imaging system (Figures 5B-E). Serial bioluminescence imaging showed generally comparable tumor signals among treatment groups on day 6 (Figures 5B-C). By day 13, mice treated with PD1-hYP218 CAR T cells showed visibly reduced tumor bioluminescent signals compared with the other treatment groups (Figures 5D-E). Tumor tissues were harvested and analyzed on days 7 and 14. The phenotypic biomarkers expressed by tumor-infiltrating hYP218 CAR T cells were compared across the different CAR T-cell groups, as well as with their respective pre-infusion CAR T-cell products via flow cytometry (Figures 5F-I, Suppl Fig. 3A-D). We observed that tumor-infiltrating PD1-hYP218 CAR T cells consistently exhibited a low PD-1 phenotype across all evaluated time points, consistent with targeted editing of the PD1 locus in these cells. In contrast, the proportion of PD-1 cells increased from approximately 20% at baseline to nearly 80% by day 14 in LV-hYP218 and AAVS1-hYP218 CAR T cells (Figures 5F, Suppl Fig. 3A). hYP218 CAR T cells across all three groups were positive for the activation marker CD69, consistent with an activated phenotype within the tumor microenvironment (Figures 5G, Suppl Fig. 3B). Non-viral hYP218 CAR T cells exhibit a higher baseline CD8:CD4 ratio (∼2.0) compared to LV CAR T cells, which have a ratio of approximately 1.1, with a transient increase in the CD8:CD4 ratio observed in PD1-hYP218 CAR T cells at day 7 (Figures 5H, Suppl Fig. 3C). Importantly, PD1-hYP218 CAR T cells maintained relatively stable expression of the CD45RA CD62L naïve phenotype over time, whereas LV-hYP218 CAR T cells exhibited a progressive decline in this population (Figures 5I, Suppl Fig. 3D). Additionally, basal expression levels of TIM3 and LAG3 were lower in non-viral hYP218 CAR T cells than in LV-hYP218 CAR T cells. The proportion of TIM3 cells decreased over time across all three CAR T-cell products, whereas LAG3 cells remained relatively stable in the non-viral hYP218 CAR T-cell groups (Suppl Fig. 4A-B). Collectively, these findings demonstrate that PD1-hYP218 CAR T cells achieved effective tumor control while maintaining a less differentiated phenotype and relatively low expression of exhaustion-associated markers in vivo.

**Figure 5.**
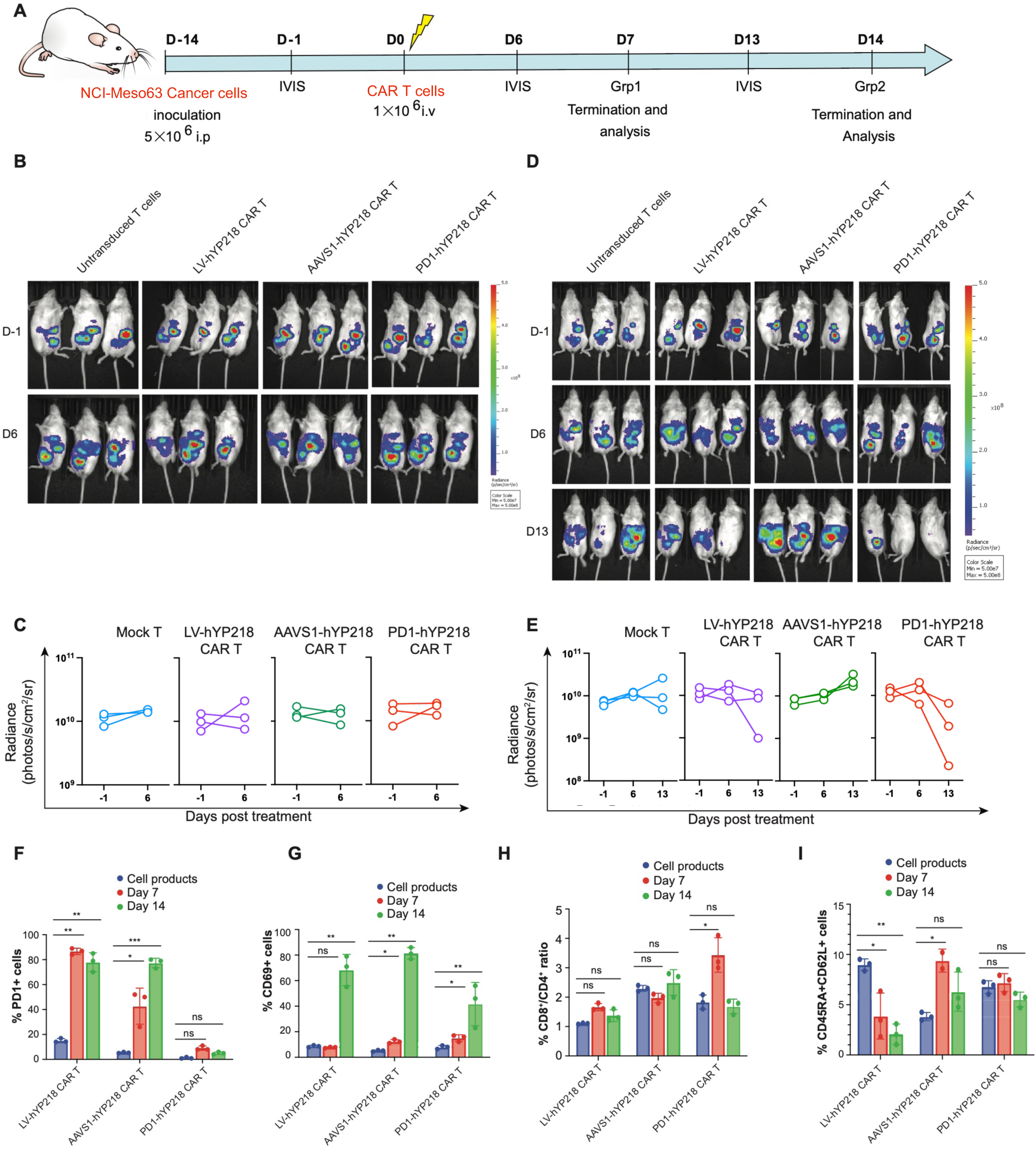
Phenotypic analysis of non-viral PD1-hYP218 CAR T-cell functionality in vivo. (A) Experimental design showing NCI-Meso63 tumor inoculation and CAR T-cell treatment. Mice bearing established NCI-Meso63 tumors were treated with 1 × 10^6^ hYP218 CAR T cells 2 weeks after tumor inoculation. (B-E) Representative BLI images and tumor burden at days 6 and 13 after treatment. B-C represent group1; D-E represent group 2. (F) PD1 expression in hYP218 CAR T-cell products and tumor-infiltrating CAR T cells at the indicated time points. (G) CD69 expression in tumor-infiltrating CAR T cells. (H) CD8/CD4 ratio. (I) Frequency of naïve (CD45RA+CD62L+) CAR T cells. Data are presented as mean ± SD (n = 3 per treatment group). ns, not significant; *P* ≤ .05; *P* ≤ .01; *P* ≤ .001.

## Discussion

Despite the remarkable success of CAR T-cell therapy in hematological malignancies, durable responses in solid tumors remain limited by persistent antigen exposure, T-cell dysfunction, and an immunosuppressive tumor microenvironment^31–33^. Site-directed non-viral genome engineering offers an opportunity to address these challenges by combining therapeutic CAR expression with modification of endogenous T-cell regulatory pathways^27,28^. Here, we developed a non-viral approach to integrate the clinically relevant hYP218 mesothelin-targeted CAR into the endogenous *PDCD1* locus and compared its functional performance with AAVS1-targeted and conventional lentiviral CAR T cells. PD1-hYP218 CAR T cells demonstrated potent antitumor activity comparable to lentiviral CAR T cells despite lower CAR surface expression, while exhibiting reduced inflammatory cytokine production and features consistent with a less differentiated T-cell phenotype.

The functional consequences of PD-1 modulation appear to depend on CAR affinity and costimulatory signaling. Previous studies have shown that HER2-targeted high-affinity CAR T cells are relatively resistant to PD-L1-mediated inhibition, while low-affinity 4-1BB-based CAR T cells exhibits resistance against PD-L1 inhibition compared with CD28-based constructs^34^. Interestingly, our study demonstrated that PDCD1-targeted integration of the 4-1BB-based hYP218 CAR supported potent antitumor activity comparable to lentiviral CAR T cells and superior to AAVS1-targeted CAR T cells, suggesting that PDCD1-targeted engineering may provide functional benefits even in CAR T cells with 4-1BB-based CAR construct. These findings raise questions about how CAR affinity, receptor density, tumor PD-L1 expression, and other features of the tumor microenvironment influence the effects of PD-1 pathway modification. Supporting the importance of tumor context, Armstrong et al. reported enhanced activity of MSLN-CAR T cells following PD-1 blockade in a PD-L1-high pancreatic patient-derived xenograft model^35^. Similarly, MUC1-directed CAR T cells engineered to secrete anti-PD-1 antibodies demonstrated improved proliferation during chronic antigen exposure and enhanced antitumor activity in cholangiocarcinoma models^36^. Together, these findings suggest that the therapeutic effects of PD-1 pathway modulation may vary with CAR design, tumor context, and the specific engineering strategy employed.

Previous studies have also linked PD-1 modulation to T-cell differentiation and functional persistence. Ouyang et al. reported that PD-1 downregulation preserved memory-associated phenotypes and reduced exhaustion-related changes in BCMA-CAR T cells^37^. More recently, Shahosseini et al. demonstrated that a PD-1/IL-15Rβ switch receptor enhanced proliferation and memory-associated features in MSLN-CAR T cells by converting inhibitory PD-1/PD-L1 interactions into IL-15-related signaling^38^. Consistent with these observations, PD1-hYP218 CAR T cells maintained a relatively stable CD45RA□CD62L□ population following infusion, suggesting retention of a less differentiated T-cell compartment during tumor exposure. Interestingly, PD1-hYP218 CAR T cells also exhibited a transient increase in the CD8□/CD4 ratio at day 7, which returned toward baseline by day 14. This early shift may reflect preferential CD8□ T-cell expansion or persistence. Furthermore, comparable TIM-3 and LAG-3 expression among CAR T-cell groups, together with the eventual decline in cytotoxicity following repeated antigen stimulation, indicate that PDCD1-targeted engineering does not fully prevent dysfunction-associated changes. Thus, the relationship between PD-1 signaling, T-cell differentiation, and functional persistence remains complex and may depend on the strength and duration of CAR-mediated activation.

A notable finding was that PD1-hYP218 CAR T cells maintained antitumor activity comparable to LV-hYP218 CAR T cells despite substantially lower CAR surface density. Although receptor abundance contributes to antigen recognition, excessive CAR expression can promote tonic signaling and T-cell dysfunction, suggesting that maximal CAR density may not confer optimal therapeutic activity^39,40^. Previous studies have shown that reduced CAR density can limit inflammatory cytokine secretion without compromising cytotoxicity^41^. Similarly, CAR T-cell products with enhanced stem-like characteristics can exhibit reduced cytokine production while maintaining cytotoxic activity^42^. Consistent with these observations, PD1-hYP218 CAR T cells produced less IFN-γ and TNF-α despite preserved tumor-cell killing, suggesting that cytotoxicity and inflammatory cytokine secretion can be partially dissociated. Together, these findings raise the possibility that CAR expression, T-cell differentiation state, and genomic integration context collectively influence therapeutic performance, although their relative contributions remain unresolved.

Emerging clinical evidence highlights both the promise and challenges of PD-1 pathway modulation in CAR T-cell therapy. Adusumilli et al. demonstrated the feasibility of combining mesothelin-targeted CAR T cells with pembrolizumab in patients with malignant pleural disease^43^, Their subsequent studies explored a CD28-based MSLN-CAR incorporating a PD-1 dominant-negative receptor and demonstrated promising preclinical activity following intraperitoneal administration^44^, although its subsequent phase I evaluation in esophagogastric cancer showed limited clinical antitumor activity despite detectable CAR T-cell persistence^45^. More recently, anti-PD-1 nanobody-armored MSLN-CAR T cells achieved an objective response rate of 63.6%, including one complete response, in patients with malignant mesothelioma^46^. Wang et al. evaluated CRISPR-engineered MSLN-CAR T cells with simultaneous PD-1 and TCR disruption in patients with MSLN-positive solid tumors, reporting limited clinical responses and reduced persistence of TCR-deficient cells^47^, suggesting that preservation of endogenous TCR signaling may contribute to CAR T-cell persistence. In comparison, our PDCD1-targeted strategy preserves endogenous TCR expression, potentially allowing additional recognition of tumor-associated antigens beyond mesothelin and providing complementary antigen-driven stimulation that may support sustained T-cell responses.

Despite the encouraging antitumor activity and distinct engineering advantages of our approach, several challenges remain before clinical translation. The approximately 10% knock-in efficiency requires further optimization for scalable manufacturing, and comprehensive assessment of off-target editing and on-target genomic alterations will be essential to establish genomic safety. Moreover, the use of immunodeficient mouse models limits evaluation of the intact tumor immune microenvironment and long-term persistence. Future studies incorporating expression-matched controls, direct assessment of PD-1-dependent signaling, and more clinically representative models will be important to distinguish the contributions of checkpoint disruption and genomic integration context and evaluate the translational potential of this approach.

In summary, our study establishes a preclinical proof of concept for non-viral integration of the hYP218 CAR into the *PDCD1* locus, supporting potent antitumor activity against mesothelin-expressing solid tumors despite relatively low CAR surface expression. With further improvements in manufacturing efficiency and genomic safety, this approach may provide a platform for developing more precisely engineered CAR T-cell therapies for solid tumors.

## Declaration

This work was supported by the Intramural Research Program of the NIH, National Cancer Institute, Center for Cancer Research. The contributions of the NIH authors are considered Works of the United States Government. The findings and conclusions presented in this paper are those of the authors and do not necessarily reflect the views of the NIH or the U.S. Department of Health and Human Services.

## Acknowledgments

We Dr. Hisataka Kobayashi and Dr. Christine Alewine of the Center for Cancer Research, National Cancer Institute, National Institutes of Health, for providing the OVCAR8 and KLM-1 cell lines, respectively.

## Author contributions

S.M., Q.J. and R.H. designed the project. S.M. performed all the experiments. R.C. assisted with plasmid design and CRISPR cloning. A.V., Y.C., and J.Z. assisted with laboratory experiments. S.M., Q.J. and R.H. analyzed the data and drafted the paper. All authors read and contributed to the final version of the paper. Conceptualization: S.M., Q.J. and R.H. Investigation: S.M., Q.J. and R.H. Methodology: S.M., A.V., R.C., Y.C., J.Z., Q.J. and R.H. Data curation: S.M., Q.J. and R.H. Validation: S.M., Q.J. and R.H. Formal analysis: S.M., Q.J. and R.H. Funding acquisition: R.H. Project administration: Q.J. and R.H. Resources: R.H. Supervision: Q.J. and R.H. Visualization: S.M., Q.J. Writing—original draft: S.M. Writing—review and editing: S.M., Q.J. and R.H. All authors read and approved the manuscript.

## Data and materials availability

All data needed to evaluate the conclusions in the paper are present in the paper and/or the Supplementary Information.

