## Supplemental Information for "Non-viral *PDCD1* locus-directed hYP218 CAR T cells with enhanced anti-tumor activity against mesothelin-positive solid tumors"

**Supplementary Information:**

**Methods and materials:**

**Construction of the hYP218 CAR cassette**

The anti-mesothelin CAR cassette is composed of the single-chain fragment variable (ScFv) derived from humanized rabbit anti-mesothelin antibody (hYP218) that binds to the membrane proximal region of mesothelin; hinge and transmembrane domains of CD8α; the 4-1BB costimulatory domain and the CD3ζ signaling domain. Transcription of the CAR element is driven by an EF1α promoter and terminated by an SV40 poly(A) signal sequence. For in vitro and in vivo tracking, as well as CAR T-cell ablation if needed to reduce toxicity, a truncated human EGFR polypeptide (huEGFRt) is added. A T2A self-cleaved sequence separates the CAR from the huEGFRt. In this study, the parental hYP218 CAR cassette is used to generate variants for integration at distinct genomic loci. The hYP218 CAR construct, intended for producing non-viral CAR T cells, is flanked by 800 bp homology arms specific for the intended integration site of the T cell genome.

**Ribonucleoprotein and linear dsDNA production**

Following activation, T cells were electroporated with ribonucleoprotein (RNP) complexes and linear hYP218 CAR DNA templates. The RNP complexes consisted of Cas9 protein and specific guide RNAs (gRNAs) targeting either the AAVS1 or PD1 locus. sgRNAs were chemically synthesized (GenScript) and resuspended in nuclease-free buffer (IDT). RNPs were produced by incubating sgRNA with recombinant spCas9 (Thermo Fisher) for 20 min at room temperature. RNPs were subjected to electroporation immediately after complex formation. For linear dsDNA production, plasmids containing the hYP218 CAR sequence flanked by 800 bp homology arms were first constructed. Linear dsDNA was then obtained by restriction endonuclease digestion and purified using the Qiagen DNA Purification kit (Qiagen). The sgRNAs used for the preparation of AAVS1 and PD1-hYP218 CAR T cells were as follows: AAVS1 sgRNA (5′-AGAGCTAGCACAGACTAGAG-3′), PD1 sgRNA (5′-CGACTGGCCAGGGCGCCTGT-3).

**PBMC isolation and activation**

PBMCs from healthy donors were provided by the NIH Clinical Center Department of Transfusion Medicine as part of its IRB-approved, consented healthy donor program. PBMC isolation from peripheral blood samples was performed using Lymphocyte Separation Medium 1077 (Promocell). Isolated PBMCs were cultured in AIM-V medium (Gibco**)** supplemented with IL-2 (200 IU/mL), IL-7 (5 ng/mL), and IL-15 (5 ng/mL) (Peprotech). Cells were pre-activated using TransAct (Miltenyi Biotech) for 48 hours at 37°C in a humidified atmosphere containing 5% CO2.

**T cell electroporation**

1×10^6^ prewashed T cells were resuspended in 20 μL electroporation buffer P3. Meanwhile, RNPs were prepared by mixing Cas9 and gRNA at a 1:1.5 ratio, followed by the addition of the DNA template (1 μg). Cells in the electroporation buffer were then added, and after proper mixing, the whole mixture was transferred to the electroporation cuvettes. Program EO115 was chosen for electroporation. After electroporation, cells were immediately supplemented with prewarmed medium, incubated for 10 minutes at 37 °C, then transferred from the electroporation cuvettes and cultured in standard 12-well plates.

**Flow cytometry**

Expression of cell-surface proteins and the CAR receptor on T cells was determined by flow cytometry. Cells were pre-washed and incubated with antibodies for 30 min at 4 °C. After washing, the samples were analyzed by flow cytometry (CytoFlex; Beckman Coulter) and analyzed using FlowJo (BD Biosciences; version 10.8.1). PE anti-human EGFR targeting the extracellular domain of huEGFRt was used as a marker for assessing hYP218 CAR expression. The percentage of hYP218 CAR+ cells was analyzed within the CD3+ gate. To evaluate PD1 knockout efficiency, flow cytometry was performed on CD3+CAR+ cells, which were stained with a fluorochrome-conjugated PD1 antibody. The percentage of PD1+ cells was compared between PD1-hYP218 CAR T cells, AAVS1-hYP218 CAR T cells, and lentiviral transduced hYP218 CAR T cells. The antibodies used for flow cytometry and immunofluorescence are included in table S1.

**hYP218 CAR** **density quantification**

Estimated CAR surface density was quantified by flow cytometry using BD Quantibrite™ PE beads (BD Biosciences). Quantibrite beads containing a defined number of phycoerythrin (PE) molecules per bead were acquired on the same day and with the same instrument settings as the CAR T-cell samples. A calibration curve was generated by plotting the median fluorescence intensity (MFI) of each bead population against the assigned number of PE molecules per bead, and linear regression was used to derive the conversion equation. For CAR T-cell samples, surface expression of the hYP218 CAR construct was assessed indirectly through staining of the co-expressed truncated human EGFR tag (huEGFRt) using a PE-conjugated anti-human EGFR antibody. Cells were gated on viable CD3⁺ lymphocytes, and the MFI of the huEGFRt-positive population was converted to PE molecules per cell using the Quantibrite calibration curve. Because huEGFRt is co-expressed with the CAR cassette through a T2A sequence, these values were used as an estimate of relative CAR surface density. CAR density was compared among LV-hYP218, AAVS1-hYP218, and PD1-hYP218 CAR T-cell products.

**Antigen restimulation assay**

The cytotoxicity and surface phenotype of LV, AAVS1, and PD1 hYP218 CAR T cells were evaluated by repeatedly stimulating them with target cells over 7 days. For this purpose, luciferase and flow cytometry assays were performed on day 0, before initiating the coculture, and on days 1, 3, 5, and 7 thereafter, for a total of 4 rounds of antigen stimulation. Briefly, hYP218 CAR T cells were cocultured with MSLN+ OVCAR8 cells at a 3:1 effector: target (E:T) ratio on day 0. After 24 hours of coculture, cytotoxicity, activation, and exhaustion-associated phenotypes were evaluated, and MSLN-CAR T cells were again restimulated with MSLN+ OVCAR8. The same procedure was repeated on days 3, 5, and 7, and at each time point, cytotoxicity and phenotype were evaluated. Cocultures were maintained for 7 days in RPMI medium without added cytokines.

**Luciferase-based cytotoxicity assays**

The Cytotoxic potential of viral and non-virally transduced hYP218 CAR T cells was assessed by coculturing them with mesothelin-expressing cancer cell lines OVCAR8 (ovarian cancer), KLM1 (pancreatic cancer), and NCI-Meso63(mesothelioma). Cytotoxicity assays were performed by plating 5000 cancer cells in 100 μL of complete RPMI medium in a 96-well plate. Four hours after seeding the cancer cells, 100 μL of culture media containing lentiviral or non-viral hYP218 CAR T cells or untransduced cells at various effector-to-target (E:T) ratios were added. For all in vitro cytotoxicity assays, equivalent numbers of CAR-positive T cells were used across all CAR T-cell products. The frequency of CAR-expressing cells was determined by flow cytometry using huEGFRt staining. After 24 hours of co-culture, cytotoxicity was assessed by measuring the luciferase signal from surviving tumor cells using the Luciferase Assay System (Promega) according to the manufacturer’s instructions.

**Cytokine profiling**

For in vitro evaluation of effector cytokine release, hYP218 CAR T cells were co-cultured

with OVCAR8, NCI-Meso63, and KLM-1 cells at an effector/target ratio of 1:1 in a regular RPMI medium without exogenous cytokines. The supernatant was collected after 24 h, and IFNγ and TNFα secretion were measured using the ELISA kit (RD Systems) according to the manufacturer’s instructions. Briefly, 100 µl of assay diluent was added to each well of the 96-well plate, followed by the addition of 100 µl of standard and samples, and the plate was incubated for 2 hours. Plates were then washed with 400 μL of wash buffer in each well, followed by the addition of the respective conjugates and incubation for 2 hours at room temperature. After incubation, wells were washed again and developed by adding the substrate solution. The reaction was stopped, and the absorbance was read at 450 nm, with background subtraction at 570 nm.

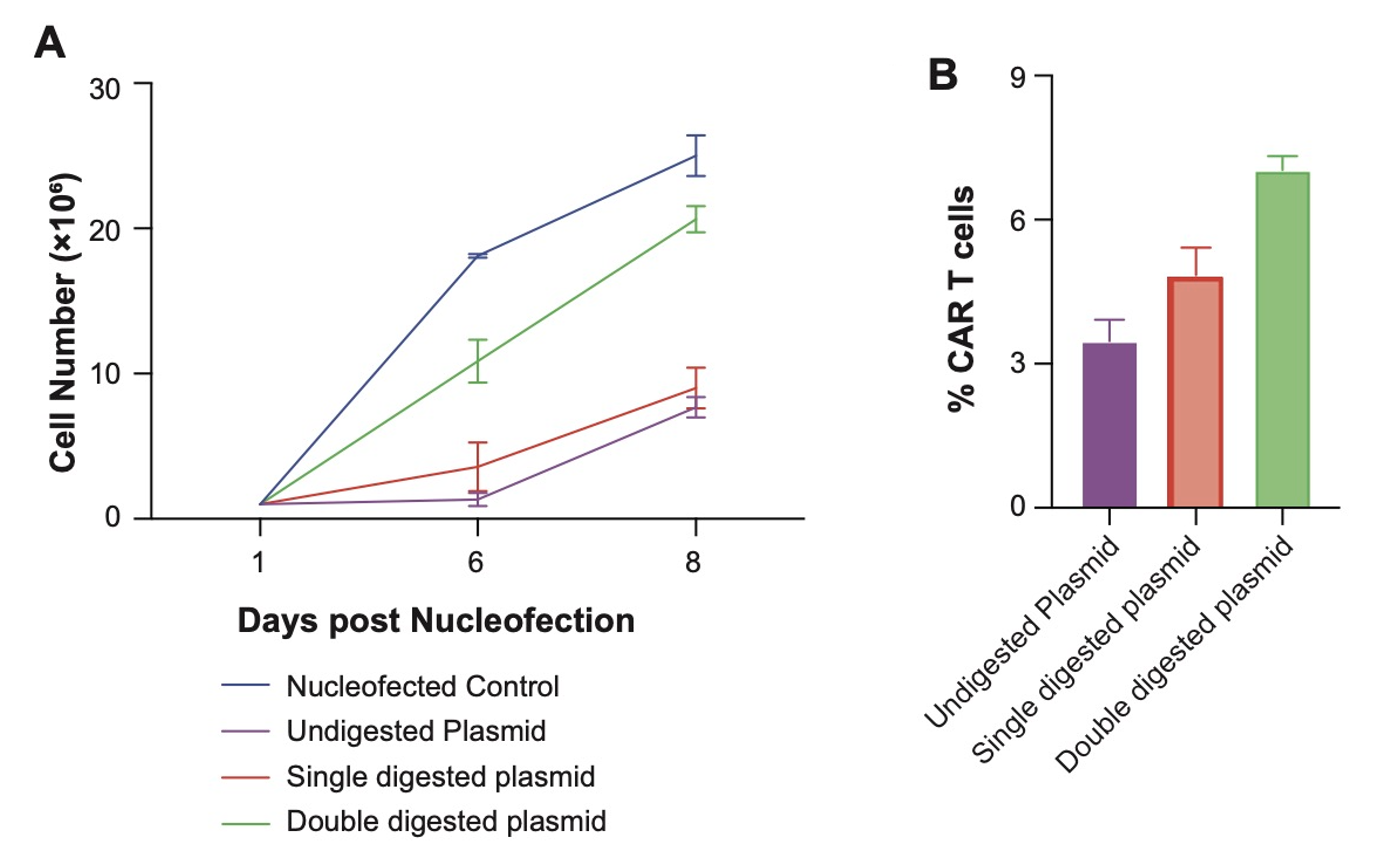

**Figure S1. Effect of plasmid linearization on PD1-hYP218 CAR T-cell production.** The PD1-hYP218 CAR plasmid (7,072 bp) was linearized by either single digestion with PciI or double digestion with DrdI and NheI. For double digestion, a 5,363-bp linear CAR donor fragment lacking bacterial propagation sequences was purified for nucleofection. (A) T-cell expansion following nucleofection with undigested plasmid or purified linearized products. T cells nucleofected with the double-digested donor exhibited greater expansion following target-cell stimulation than those nucleofected with undigested or single-digested plasmids. (B) CAR expression, determined by flow cytometric detection of EGFR-positive T cells, was higher following nucleofection with the double-digested donor than with undigested or single-digested plasmids. A total of 0.6 µg DNA was used per nucleofection.

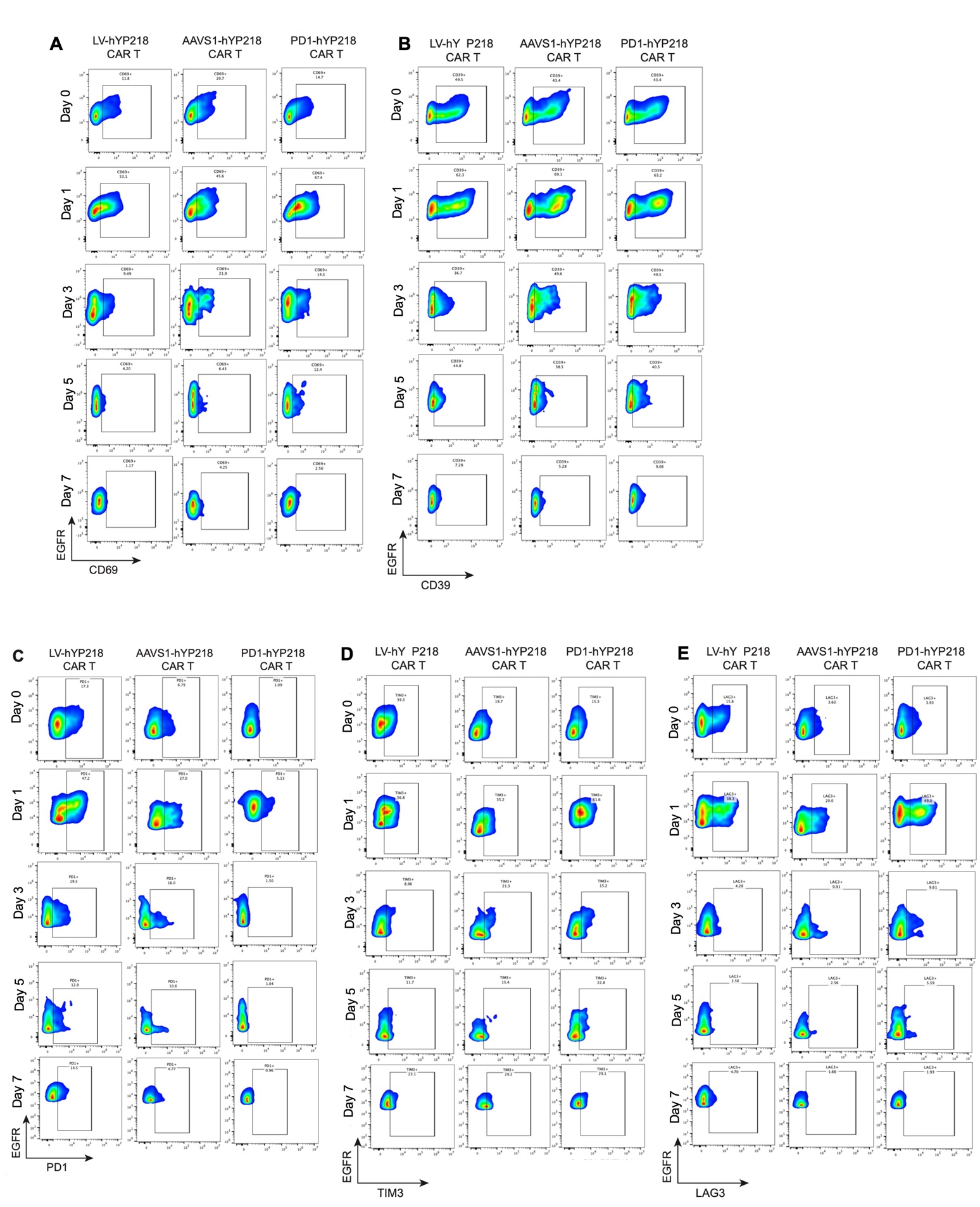

**Figure S2. Phenotypic analysis of non-viral PD1-hYP218 CAR T-cell in vitro**. CAR T Cells were in vitro stimulated with tumor cells for four times and harvested at different time points during the 7-day coculture. The indicated biomarker expression in CAR T cell was detected by flow cytometry, including CD69 (A), CD39 (B), PD1 (C), TIM3 (D), LAG3 (E).

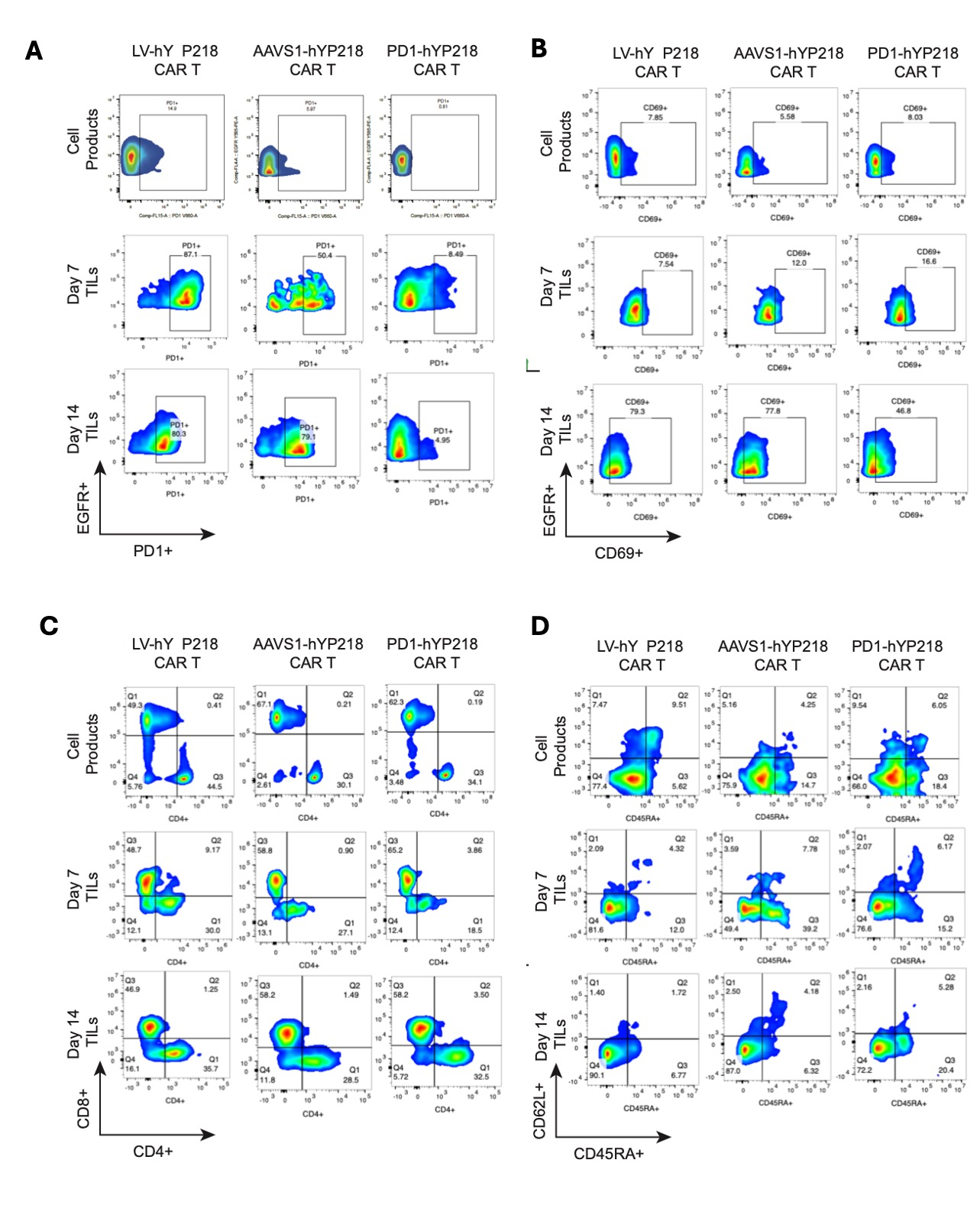

**Figure S3**. **Phenotypic analysis of non-viral PD1-hYP218 CAR T-cell in vivo.**

Representative biomarkers expressed by PD1-hYP218 CAR T detected by flow cytometry. Mice bearing established NCI-Meso63 tumors were treated with 1 × 10^6^ hYP218 CAR T cells. Tumor tissues were harvested at indicated time points for analysis. (A) PD1 expression in hYP218 CAR T-cell products and tumor-infiltrating CAR T cells at the indicated time points. (B) CD69 expression in tumor-infiltrating CAR T cells. (C) CD8 ratio. (D) Frequency of naïve (CD45RA+CD62L+) CAR T cells.

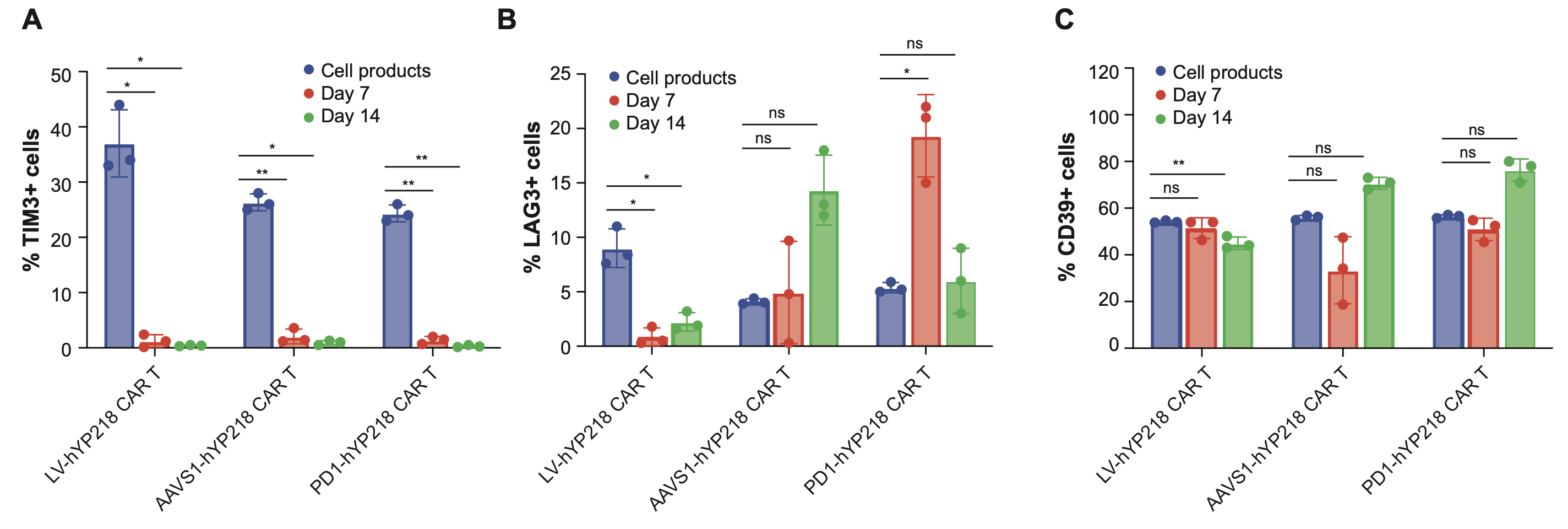

**Figure S4.** (A) Comparison of the percentage of TIM3+ tumor-infiltrating LV, AAVS1, and PD1-hYP218 CAR T cells, evaluated at Day 7 and Day 14 post-infusion. A significant decrease in the percentage of TIM3+ cells was observed over time for all the hYP218 CAR T cell products, compared to the initial cell product baseline. (B) A significant decrease in the percentage of LAG3+ cells was observed over time in LV hYP218 CAR T cells. non-viral hYP218 CAR T cells, maintain LAG3+ phenotype. (C) LV hYP218 CAR T cells show decrease in CD39% cells by day 14, while as an increase in the CD39 was observed at day 14 in PD1-hYP218 CAR T cells. Error bars represent the standard deviation (SD) of the measurements. Statistical significance was determined using unpaired Student's tests (*p < 0.05, **p < 0.01).

**Supplementary Table 1. Antibodies used in Flow cytometry**

| **REAGENT or RESOURCE** | **SOURCE** | **IDENTIFIER** |
| --- | --- | --- |
| CD62L Antibody, anti-human PE-Vio@770 | MACS Miltenyi Biotec | 130-113-621 |
| IL15 Monoclonal Antibody | Invitrogen | MA5-23561 |
| PE anti-human IL21 | BioLegend | 513003 |
| PE/Cyanine7 anti-human | BioLegend | 372714 |
| APC anti-human CD45RA | BioLegend | 304150 |
| APC anti-human CD3 | BioLegend | 300312 |
| BV650 Mouse Anti-human CD279 (PD-1) | BD Biosciences | 564104 |
| APC/cyanine7 anti-human CD25 | BioLegend | 302614 |
| anti-hTGF-b R11 | R&D Systems | FAB241F |
| FITC anti-human CD8 | BioLegend | 344704 |
| FITC anti-human CD274 | BioLegend | 393606 |
| APC anti-human CD274 | BioLegend | 329708 |
| Brilliant Violet 785 anti-human LAG-3 | BioLegend | 369322 |
| Brilliant Violet 785 anti-human CD69 | BioLegend | 310932 |
| APC anti-human CD4 | BioLegend | 300514 |
| FITC anti-human CD39 | BioLegend | 328206 |
| Brilliant Violet 510 anti-human (Tim-3) | BioLegend | 345030 |
| Rabbit anti-EGFR | Novus a Biotechne Brand | NBP2-52671 APC |
| Brilliant Violet 510 Mouse | BioLegend | 400171 |
| R718 Mouse IgG2a,k isotype control | BD Biosciences | 566928 |
| FITC Mouse IgG2a,k isotype control | BD Biosciences | 554647 |
| PE Mouse IgG2b, k isotype | BioLegend | 400314 |
| APC Mouse IgG1 | BioLegend | 400120 |
| Brilliant Violet 785 Mouse IgG1 | BioLegend | 400169 |
| PE/CY7 Mouse IgG2a | BioLegend | 400253 |
| BrilliantViolet 510 Mouse IgC1 | BioLegend | 400171 |
| BV 650 Mouse IgG,k | BD Biosciences | 563231 |
